# Three-dimensional beating dynamics of *Chlamydomonas* flagella

**DOI:** 10.1101/2020.07.20.212159

**Authors:** Soheil Mojiri, Sebastian Isbaner, Steffen Mühle, Hongje Jang, Albert Johann Bae, Ingo Gregor, Azam Gholami, Jörg Enderlein

**Author notes:** **For correspondence:** (AG); (JE). **Present address:**Department of Biomedical Engineering, University of Rochester, USA.

## Abstract

Axonemes are the basic structure of motile cilia and flagella, and the investigation of how they function and move requires rapid three-dimensional imaging. We built a multi-plane phase-contrast microscope for imaging the three-dimensional motion of unlabeled flagella of the model organism *Chlamydomonas reinhardtii* with sub-μm spatial and 4 ms temporal resolution. This allows us to observe not only bending but also the three-dimensional torsional dynamics of these small structures. We observe that flagella swim counter-clockwise close to a surface, with negatively-valued torsion at their basal and positively-valued torsion at their distal tips. To explain the torsional dynamics and signature, we suggest the existence of an intrinsic negative twist at the basal end that is untwisted by active positive-twist-inducing dynein motor proteins. Moreover, dyneins walking towards the basal induce an opposite twist at the distal tip. Bending of the whole axoneme structure then translates this twist into an observable torsion. This interconnection between chiral structure, twist, curvature, and torsion is fundamental for understanding flagellar mechanics.

---

Flagellar and ciliary motion is fundamentally important for life. Flagella of single-celled organisms are required for locomotion in fluids and thus photo- and chemotaxis. Flagella drive sperm cells to the egg for fertilization. Carpets of cilia on the respiratory epithelium are life-important for mucociliary clearance of the airways of large animals or for active fluid transport in the brain Wanner et al. (1996); Brooks and Wallingford (2014); Pellicciotta et al. (2019); Faubel et al. (2016). Motion of nodal cilia is necessary for correct embryo development Hirokawa et al. (2006); Satir and Christensen (2007); Smith et al. (2019). Thus, understanding the mechanism and dynamics of motile flagella and cilia is of paramount importance. Their cytoskeletal core, the axoneme, is a highly conserved structure. It consists of nine microtubule doublets (MTDs) forming a cylinder that surrounds a tenth pair of microtubules on its axis, with the outer MTDs being decorated with dynein motor proteins. Both the radial and longitudinal polarities of the axonemal structure Bui et al. (2012) introduce an inherent chirality of its structure. This can be seen by the orientation of the dynein-decorated MTDs around the central MTD. Fueled by hydrolyzation of adenosine triphosphate (ATP), asymmetric activity of dyneins Brokaw and Kamiya (1987); Bui et al. (2012); Lin and Nicastro (2018) slides neighboring MTDs with respect to each other which drives a regular motion in an intact axoneme Brokaw (1989); Sanchez et al. (2011).

One of the most prominent examples of a flagellum-based motile system is the single cellular bi-flagellate green alga *Chlamydomonas reinhardtii* which swims in a breaststroke manner using both stochastic and in-phase flagellar beats Goldstein et al. (2009); Polin et al. (2009); Wan et al. (2014). The grown flagella of *Chlamydomonas* have a quite stable length between 10 and 12 μm Hendel et al. (2018); Orbach and Howard (2019), which has recently been shown to be the optimum length for efficient mechanical beating Bottier et al. (2019). *Chlamydomonas* has been extensively studied as a biological model system to elucidate flagella-driven propulsion Ringo (1967); Brokaw and Luck (1983); Guasto et al. (2010). Detachment and isolation of flagella from the *Chlamydomonas* cell body Craige et al. (2013); Witman (1986) tremendously facilitates the study of their structure and dynamics, from the level of a full axoneme down to individual microtubules. In particular, it allows to investigate the mechanism of pure axonemal motion in a highly controlled and reproducible manner Bessen et al. (1980); Geyer (2013).

In this work, we study the motility of isolated wild-type reactivated axonemes of *Chlamydomonas.* The majority of published literature considers Chlamydomonas flagellar beating and the mechanisms behind purely in two dimensions Brokaw (1971); Witman et al. (1978); Lindemann (1994); Riedel-Kruse et al. (2007); Mitchison and Mitchison (2010); Sartori et al. (2016b); Lin and Nicastro (2018). Although observation of non-planar flagellar motion has been reported for various micro-swimmers such as human sperm Ishijima et al. (1986); Gadêlha et al. (2019); Su et al. (2012), bull sperm Merola et al. (2013); Daloglu and Ozcan (2017), sea urchin sperm Gong et al. (2020); Jikeli et al. (2015), the alga *Euglena’s* flagellum Rossi et al. (2017), or *P. berghei* microgametes Wilson et al. (2013), nearly all experimental studies of *Chlamydomonas* axoneme motility were purely two-dimensional. This is due to the fact that Chlamydomonas axonemes are almost three to five times shorter and faster (~50 Hz beat frequency) than flagella of typical sperm cells. Therefore, real-time 3D investigation of flagellar motion requires a rapid imaging technique with high spatial resolution.

Here, we present a novel multi-plane phase-contrast microscope that combines a recently developed multi-plane beam-splitter Geissbuehler et al. (2014); Descloux et al. (2018) with a Zernike phase-contrast optics to enable rapid, label-free, robust and easy-to-use 3D imaging of flagellar motion. Our multi-plane phase-contrast microscope allows for the simultaneous recording of eight axially stacked images with several hundred frames per second. In contrast to quantitative phase imaging or light-field imaging, our system does not require any reference image for image reconstruction, and it provides real-time 3D observation of rapidly moving samples with excellent signal-to-noise ratio (SNR) in a robust and straightforward manner.

## Results

A schematic of our new custom-built multi-plane phase-contrast microscope is shown in Fig. 1 (for details see Methods section and Appendix 1). The system allows for parallel recording of eight wide-field images in eight focal planes that are evenly spaced along the optical axis (inter-plane distance 335 nm). Using this system, we recorded movies of beating axonemes with 272 volumes per second. An example of four of such consecutively taken volumetric images are shown in Fig. 2a, where the color encodes the third dimension. In each of the volumetric images, the three-dimensional contour of an axoneme was discretized, and the resulting discrete lateral *X_j_* and *y_j_* and axial *z_j_* positions were fitted with polynomial functions (for details see Methods section). The result is a four-dimensional representation **r**(*s,t*) of an axoneme’s contour as a function of time *t* and arc length *s*, where we have |*∂***r**(*s,t*)/*∂s*| ≡ |**r′**(*s,t*)| = 1. Discretized contours and fit results for the four images in Fig. 2a are presented in Fig. 2b. As seen from the projections into the different coordinate planes, the discretized axial positions show a ~10-fold bigger jitters than the lateral positions. We estimated the accuracy of the polynomial contour fit from the standard deviations between fitted and discretized position values, and find a lateral position accuracy better than 20 nm, and an axial position accuracy better than 120 nm (see Fig. 12 in Appendix 4).

**Figure 1.**
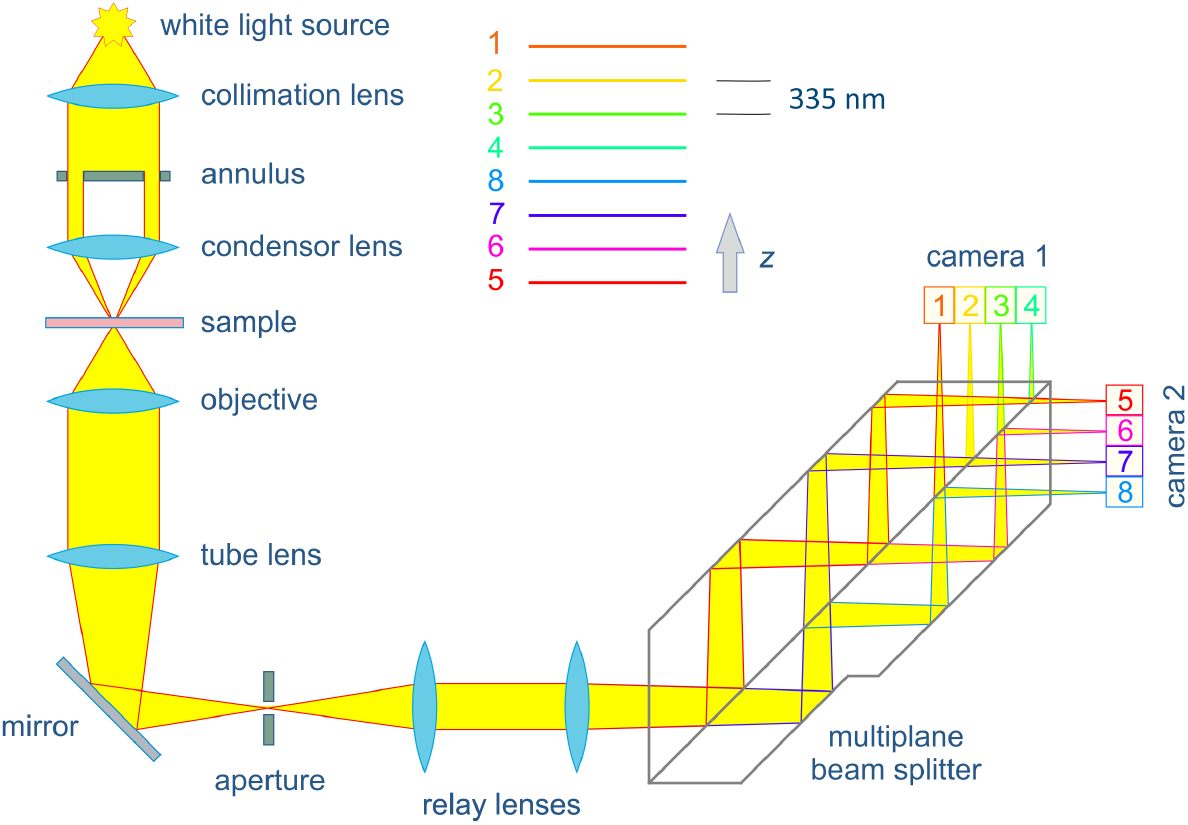
Schematic of the multi-plane phase-contrast microscope. Light from a white-light source is sent through an annular aperture and focused onto the sample for high-angle illumination. Transmitted and scattered light are collected with a Zernike phase-contrast objective (oil immersion, 60× magnification, 1.25 NA), focused by the tube lens through a slit aperture, and then sent through two relay lenses (that increase magnification to 80×) towards a custom-made multi-plane beam splitter. This splitter generates eight laterally shifted image replicas with increasingly longer optical path length, corresponding to eight different focal planes in the sample. These eight images correspond to eight focal planes in the sample separated by 335 nm (in aqueous solution), and they are recorded by two sCMOS cameras. The slit aperture limits the field-of-view and prevents overlap between neighboring images on the cameras.

**Figure 2.**
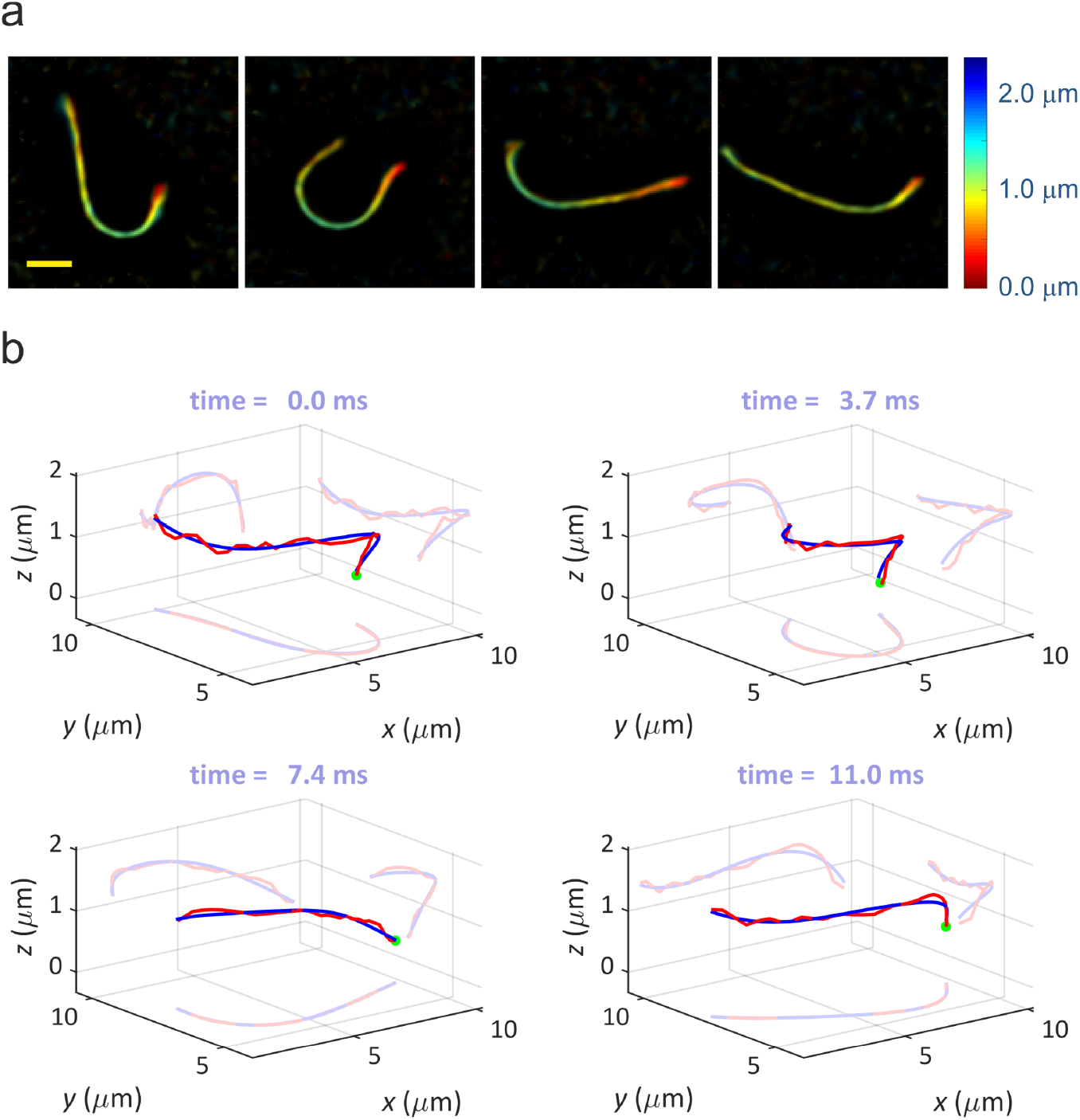
Three-dimensional recording of axoneme motion. (a) Four consecutive raw images recorded with the multi-plane phase-contrast microscope. Color encodes the axial position (see color bar), the yellow bar in the leftmost panel has a length of 2 μm. (b) Discretized axoneme contours (red) and fitted polynomial curves (blue) for the four images shown in (a). For better visibility, the respective projections into the three coordinate planes are also shown. The green dot indicates the basal end of the axoneme.

From the polynomial representation of the axonemal contours, spatio-temporal profiles of curvature *κ*(*s,t*) and torsion *τ*(*s,t*) are calculated by Do Carmo (2016)

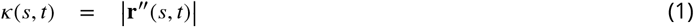

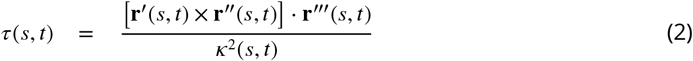

where a prime denotes differentiation with respect to arc length *s*. It should be emphasized that torsion quantifies the out-of-plane bending of an axoneme and can only be extracted from its full three-dimensional contour.

We measured the contour dynamics of 35 axonemes in a normal aqueous-like swimming medium. All observed axonemes move close and parallel to the coverslide surface of the observation chamber, due to hydrodynamic interactions with the surface Lighthill (1976); Elgeti et al. (2010). All except one of them showed a counter-clockwise (ccw) circular motion (when viewed from above). For further analysis, here we present two ccw moving flagella with nearly identical swimming radius and beat frequency (~73 Hz, determined from Fourier power spectra of space-time curvature plots). Projections (*x,y*-coordinates) of the swimming contours of the basal end and distal tip for both flagella are shown in Figs. 3a and 3c. The basal end traces a smaller circle and swims closer to the surface than the distal end and clearly follows a helical path (see Fig. 2b and supplemental movies ‘Video1’ and Video2’).

**Figure 3.**
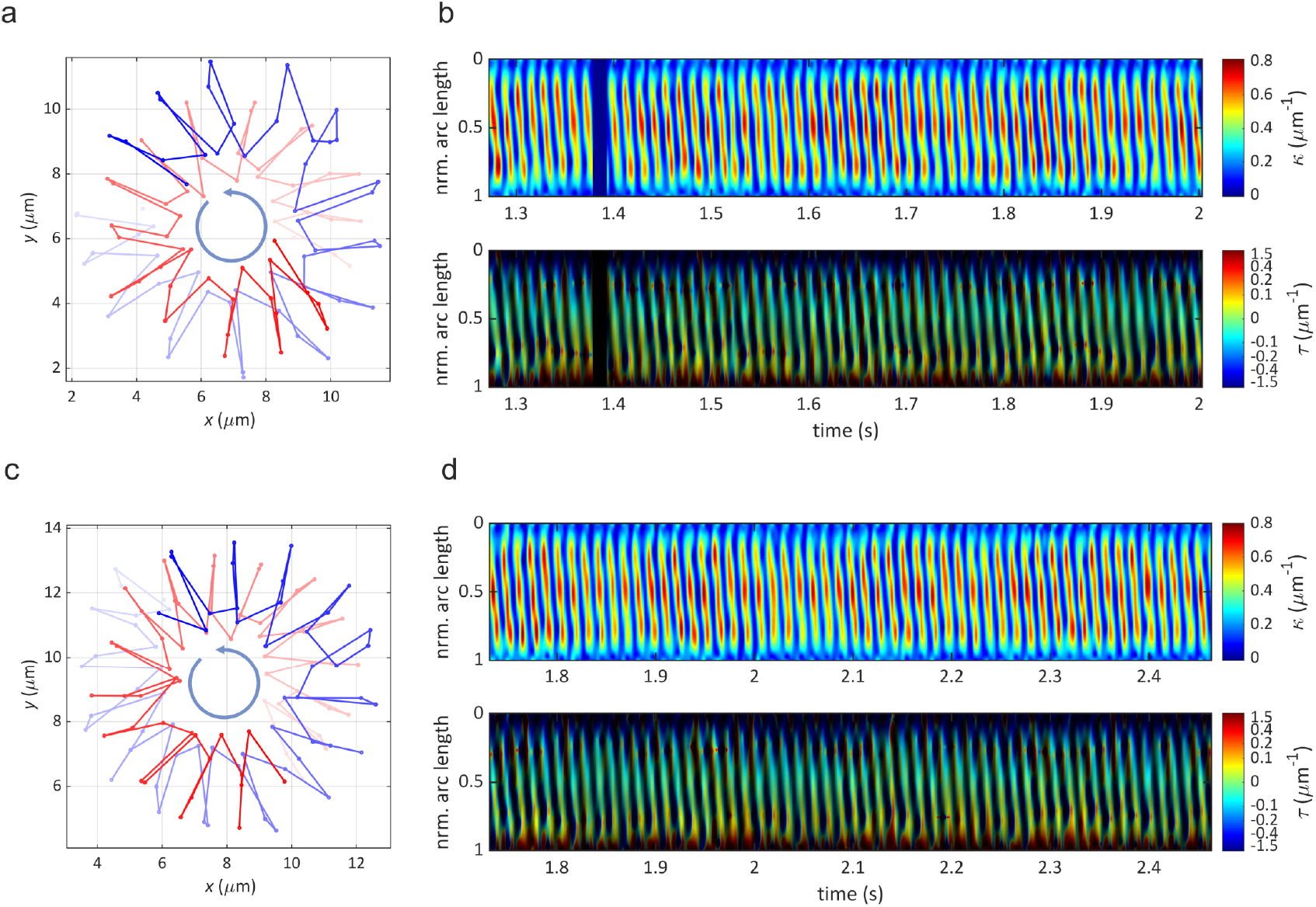
Analysis of flagellar motion. (a,c) Circular motion of basal end (red) and distal (blue) tip of two ccw-moving flagella as see from above the surface. From light to dark shading indicates progression in time. Time between two discrete points of trajectory is 3.7 ms. (b,d) Space-time plots of curvature and torsion for the same flagella as shown in (a,c), over an observation time of ~ 0.7 s. The dark vertical lane at ~1.4 s corresponds to a short excursion of the distal tip of the flagellum beyond the axial range of our observation volume and was excluded from analysis. Vertical axis is the *normalized* arc length (*s* = 0 at basal end, *s* = 1 at distal tip). In the torsion plots, color encodes torsion values, while brightness is proportional to curvature. This suppresses nonphysical torsion values in regions of low or zero curvature, where torsion is ill-defìned. Note also the strongly non-linear color mapping (see color bar) which is used to make small torsion values better visible.

Space-time plots of curvature and torsion for both flagella are presented in Figs. 3b and 3d (see supplemental movies ‘Video1’ and ‘Video2’ for the corresponding three-dimensional flagellar motion). The plots show curvature waves that start at the basal end (*s* = 0) and move towards the distal tip. This is accompanied by torsion waves that start with negative torsion values at the basal end and gradually change sign until they finish with positive torsion values at the distal tip. This torsion wave dynamics is nearly identical for both the presented axonemes.

The observed bending and torsion dynamics demonstrates the fundamentally *three-dimensional* nature of flagellar motion. We propose that the observed torsional waves are closely connected to the intrinsic chiral structure of the axoneme and to twist induced by dyneins. According to the model developed in Refs. Brokaw (2002) and Sartori et al. (2016a), dynein motors induce sliding between neighboring MTDs which does not only drive axoneme bending but also induces twisting of the axoneme. Dyneins transiently attach with their MT binding domains to neighboring MTDs and walk towards the basal end which cause distal-oriented sliding forces between neighboring MTDs Lin et al. (2014). In an untwisted axoneme, such a motion will induce a positive twist (as seen from the basal end) which should be observable as a positive torsion in a bend axoneme. Because we observe a repetitive negative torsion in each beat cycle close to the basal end we suggest the existence of an intrinsic negative twist at the beginning of each beat cycle. This is visualized in Fig. 4a as a side view of sinistrally twisted MTDs close the basal plate of an axoneme to which the axonemal MTDs are fixed (note that numbering of MTDs in this figure follows Ref. Dutcher (2020)). Then, action of the dyneins unwinds this negative intrinsic twist, reducing negative torsion at the basal end but generating positive twist and torsion at the initially twist- and torsion free distal end (Fig. 4b). Such change in torsion sign has been observed also in quail sperm flagella Woolley and Vernon (1999). Any bending of the axoneme will automatically translates twist into torsion with the same sign.

**Figure 4.**
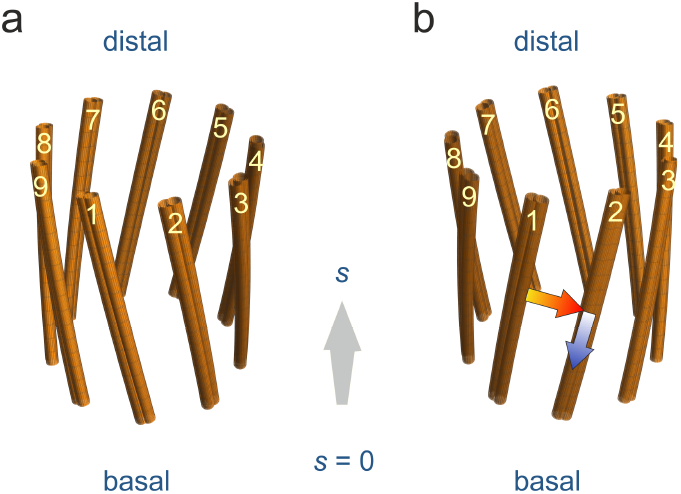
Twist of an axonemal structure. Shown are the nine MTDs around the circumference of an axoneme. (a) Intrinsic sinistral twist close to the basal end. (b) dynein-induced dextral twist close to the distal tip. The red arrow indicates the direction along which dynein motors reach from their MTD of attachment to the neighboring MTD on which they act. The blue arrows indicate the walking direction of the dynein motors. The action of the dyneins leads to sliding of MTDs with respect to each other. Due to structural connections between MTDs (e.g. by nexin proteins), this is likely to induce local twists of the axoneme structure, as shown in 4(b). If the whole structure is additionally bent, this twist would translate into torsion (with corresponding to twist sign), as experimentally observed at the basal end and distal tip of the bare axonemes.

## Discussion and Conclusion

We have developed a novel multi-plane phase-contrast microscope that allows for volumetric imaging with close to diffraction-limited resolution and with rates of several hundred volumes per second. In our application of imaging *Chlamydomonas* axonemes we have chosen such a magnification so that the eight image planes cover a volume of ~2.5 μm depth. For applications that require imaging over a deeper volume, this depth value can be easily changed by changing magnification. Using our microscope, we resolved the rapid ATP-driven motion of demembranated *Chlamydomonas* flagella in three dimensions. We were able to reconstruct the spatio-temporal dynamics of curvature and torsion of axonemes and proposed that the observed torsional waves are due to dynein-generated axoneme twisting determined by axoneme structural chirality and dynein walking direction. Torsion waves start with a negative sign at the basal end and gradually change sign while propagating towards the distal tip. We attribute the negative sign of torsion to a left-handed twist at the proximal side which slowly switches to a right-handed twist as we move towards the distal tip. Since dyneins are minus-directed molecular motors, active torque generation of accumulated dyneins at the distal tip of axonemes can generate a right-handed twist, consistent with the sign of the positive torsion observed at this location. We hypothesize that the left-handed twist at the basal end is a passive structural twist that exists independent of activity of dynein motors. In future studies, it would be highly interesting to see whether this connection between torsion dynamics and structural twist can be corroborated, for example by electron-microscopy of flagella. Also, which type of dyneins contribute to non-planar motion requires further investigations.

## Methods

### Multi-plane phase-contrast microscope

Our custom-built multi-plane phase-contrast imaging system (Fig. 1) is established on a commercial IX71 microscope from Olympus. The collimated light of a white-light source (Halogen Lamp U-LH100L-3) is send through a condenser annulus (IX-PH3, Olympus) and focused by a condensor lens (IX2-LWUCD, Olympus) onto the sample (Köhler illumination). Scattered and transmitted light are collected by a Zernike phase contrast objective (UPLFLN 60XOIPH, Olympus, 1.25 NA, 60× magnification) which shifts the phase of the transmitted light by-π/2 with respect to the scattered light resulting in destructive interference between scattered and non-scattered light. This provides a phase contrast image when focused by the tube lens onto a camera. Before imaging, a custom-made multiple beam-splitter prism splits the collected light into eight beams with different optical path lengths corresponding to eight different focal planes in the sample (see section 6.2 in the SI of Ref. Descloux et al. (2018) for a detailed explanation of the prism design). These focal planes are equally spaced with a distance of ~335 nm along the optical axis, thus spanning a total volume of ~2.4 μm depth (see Fig. 1 in Appendix 1 for calibration of the distances between focal planes). Four of the light beams exit the beam-splitter prism in a direction parallel to the incoming light, and four exit the prism in a perpendicular direction. Each set of these four beams is imaged onto a separate camera in such a way that the images of the different focal planes are positioned next to each other on the camera chip (see Fig. 1a,b in Appendix 2). The rectangular field stop aperture in the focal plane of the tube lens adjusts the field of view and to prevent overlap between adjacent images on the cameras. Since the geometry and refractive index of the prism is fixed, the inter-plane distance can be changed only by changing the magnification of the imaging system (see Appendix 1 for’setup calibration’). Image acquisition by the two cameras is synchronized with an external trigger. Data acquisition is controlled by the open-source microscopy software Micro-Manager (https://micro-manager.org/). Table 1 lists the specifications of all optical components of the setup. A detailed procedure of data processing before 3D axoneme tracking is explained in Appendix 2.

### Axoneme isolation and reactivation

Axonemes are isolated from wild-type *Chlamydomonas reinhardtii* cells, strain SAG 11-32b. They were grown axenically in TAP (tris-acetate-phosphate) medium on a 12 h/12 h day-night cycle. Flagella were isolated using dibucaine Witman (1986); Alper et al. (2013), then purified on a 25% sucrose cushion, and demembranated in HMDEK(5 mM MgSO_4_, 30 mM HEPES-KOH, 1 mM EGTA, 50 mM potassium acetate, 1 mM DTT, pH 7.4) supplemented 0.2 mM Pefabloc. The membrane-free axonemes were re-suspended in HMDEK plus 1% (w/v) polyethylene glycol (m_w_ = 20 kg/mol), 30% sucrose, 0.2 mM Pefabloc and stored at −80°C. To perform reactivation experiments, axonemes were thawed at room temperature, then kept on ice and were used for up to 1 hr. Thawed axonemes were diluted in HMDEKP reactivation buffer containing 1 mM ATP and infused into 100 μm deep flow chambers, built from cleaned glass and double-sided tape. The glass surface was blocked using casein solution (from bovine milk, 2 mg/mL) to avoid attachment of axonemes to the substrate.

### Axoneme contour tracking

Tracking and contour fitting for observed axonemes is split into two parts: In a first step, a 2D maximum projection image (along the optical axis) is generated. Tracking of this 2D image is done using a gradient vector flow snake Xu and Prince (1997); Xu et al. (1998). In a second step, the axial position (*z*) of each segment is found by fitting a 1D Gaussian to its intensity profile along the *z*-axis at fixed (*x,y*)-positions (see Appendix 3). The tracking routine yields a discrete approximation of an axoneme’s contour represented by a set of 30 three-dimensional positions **r***i* = (*x_i_, y_i_, z_i_*), *i* = 1…30. For each time frame, these points constitute a polygonal chain, which is least-square fitted with polynomial functions of the form 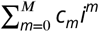, where the *c_m_* are fit coefficients. We have set *M* = 6 (6^th^ order polynomial) for fitting the lateral *x,y*-positions, and *M* = 3 (3^rd^ order polynomial) for the axial *z*-positions.

To suppress nonphysically large curvature values at the contour ends, we regularized the contour by extending it on both ends with short mirror-symmetric patches before polynomial fitting. More precisely, at the *i* = 1 and, we add additional points via **r**_1 −*j*_ = 2**r**_1_ − **r**_1 +*j*_, and at the *i* = 30 end, via **r**_30 + *j*_ = 2**r**_30_ − **r**_30 − *j*_. This changes the absolute curvature values at the ends of a flagellum, but impacts only slightly the results in the middle (see Fig. 1 in Appendix 4). For our final analysis, we have chosen a padding with three additional points on each end.

## Acknowledgment

SH thanks the European Research Council for financial support of his position via the International Trainee Network (ITN) BE-OPTICAL (grant number 675512). SM and JE acknowledge financial support by the Deutsche Forschungsgemeinschaft (DFG, German Research Council) via the Collaborative Research Center SFB 937 “Collective behavior of soft and biological matter,” project A11. AG and AB acknowledge support by MaxSynBio Consortium, which is jointly funded by the Federal Ministry of Education and Research of Germany and the Max Planck Society. A.G. thanks M. Lorenz and the Göttingen Algae Culture Collection (SAG) for providing the *Chlamydomonas reinhardtii* strain SAG 11-32b.

## Appendix 1 Setup calibration

The multi-plane beam-splitter (Fig. 1) splits the collected light into eight channels which are imaged in parallel on two cameras. The splitter was designed to render four adjacent channels with interleaved lateral distance of *d* = 3.2 mm in between which fits in size to width of the 13.3 × 13.3 mm pixel area of our sCMOS detector (ORCA-Flash 4.0 V2, Hamamatsu). Regarding focus of the camera 1 as the reference, the camera 2 is displaced from the focus by a distance of 4*d/n* where *n* = 1.46 is the refractive index of the prism material. In other words, with respect to the prism outputs, camera 2 is positioned by distance of 4*d/n* more away than the camera 1. This allows one to obtain an optical path difference of ~*d/n* between sequential planes. To calibrate this system, we performed *Z*-scans of immobilized green fluorescent beads of 200 nm diameter (FluoSpheres− Carboxylate Microspheres, 0.2 μm, yellow green fluorescent (505/515), Thermo Fisher Scientific) that were spin-coated on a glass coverslide, using an oil immersion phase objective (60×, 1.25 NA) and the relay lens system shown in Fig. 1. Fluorescence excitation was done at 470 nm with a LED, and the objective was moved over an axial scan range of6 μm with 100 nm step size. Fig. 1a displays multi-plane image acquisition of fluorescent beads at one *z*-scan step.

**Appendix 1 Figure 1.**
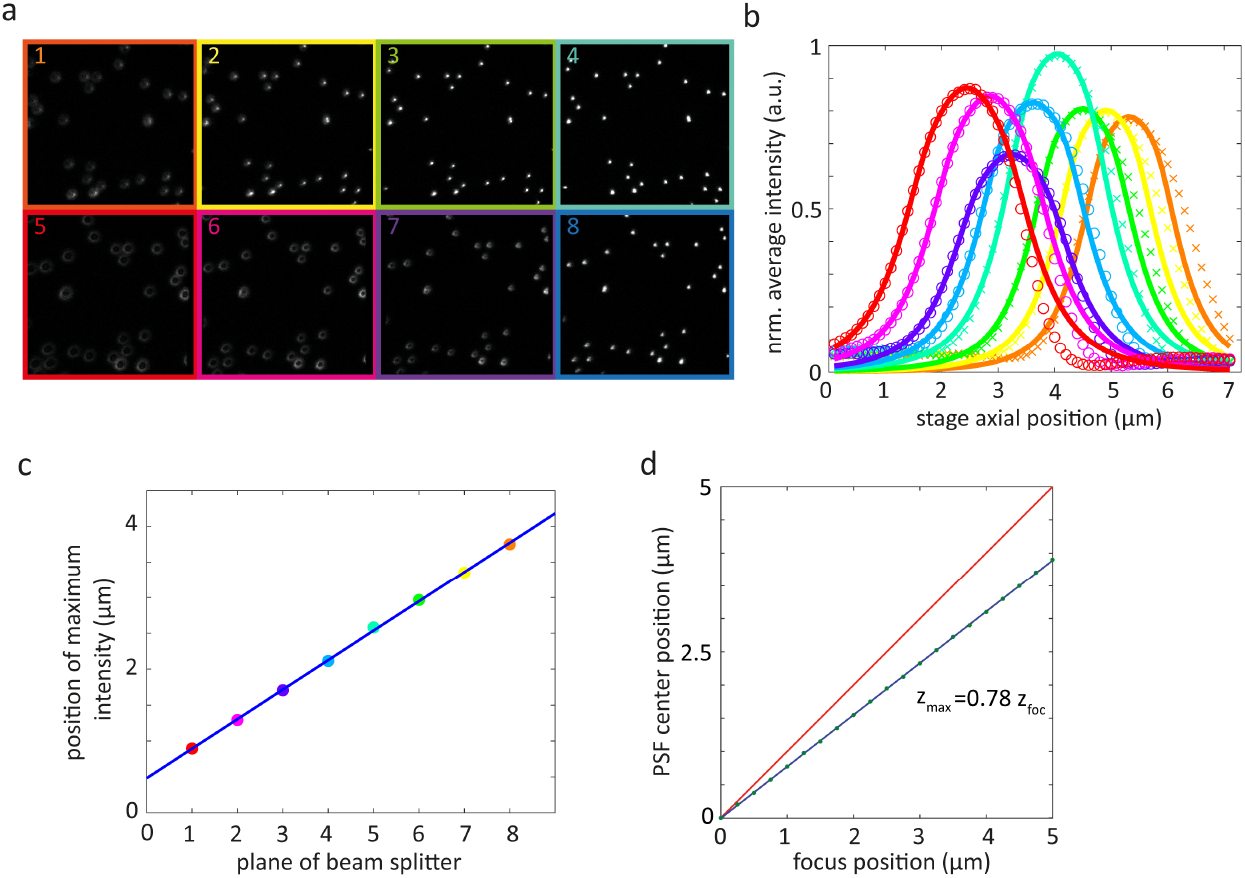
Inter-plane distance and brightness calibration. (a) Example of the eight images of fluorescent beads recorded by the multi-plane wide-field microscope. In this example, beads are in focus in nominal plane #4 (b) Normalized average intensity in each image plane as a function of axial scan position (for axial scan steps of 100 nm). Crosses (for camera 1) and open circles (for camera 2) represent measured data, solid lines are Gaussian fits. (c) Linear fit of positions of intensity maxima from panel (b). The fit yields an average inter-plane distance of ~430 nm (for oil immersion). (d) Relative shift of focal plane position in water (blue line) with respect to oil (red line). Open circles are the result of a wave-optical calculation of imaging in water, the blue solid line is a linear fit to this result. It shows that close to the glass interface, we still find a linear relationship between focal plane position and the objective’s axial position, but inter-plane distance in water is by a factor of 0.78 smaller than in oil.

A plot of the total intensity recorded in each of the eight planes as a function of axial position of the objective together with Gaussian fits is shown in Fig. 1b. A linear fit of the position of their maxima as a function of axial position of the objective, Fig. 1c, yields an average inter-plane distance of ~430 nm. The corresponding inter-plane distance in water shrinks by a factor of 0.78 due to the refractive index mismatch between immersion oil and water, as numerically calculated using the full wave-optical theory developed by Wolf and Richards Wolf (1959); Richards and Wolf (1959). The result of these calculations, Fig. 1d, shows how the axial center of the PSF in water (blue line) deviates from its position it would have in glass/oil (red line). Thus, our multi-plane system allows us to record a sample volume of ~40 × 40 × 2.4 μm in an aqueous medium.

The area under each curve in Fig. 1b is the relative amount of light that is recorded in each of the eight channels. These numbers, ε(plane #), are required later for correcting the slight brightness variations between different planes that originate from a non-perfect equal division of light by the beam splitter (±11.7% s.d. of averaged brightness).

## Appendix 2 Work flow from raw data to registered 3D image stack

### Image acquisition

Images in eight focal planes, divided into two groups of four upper and four lower planes, are simultaneously acquired by camera 1 and camera 2, respectively (Fig. 1a,b). Images of camera 1 are horizontally flipped to have the same orientation as those of camera 2.

**Appendix 2 Figure 1.**
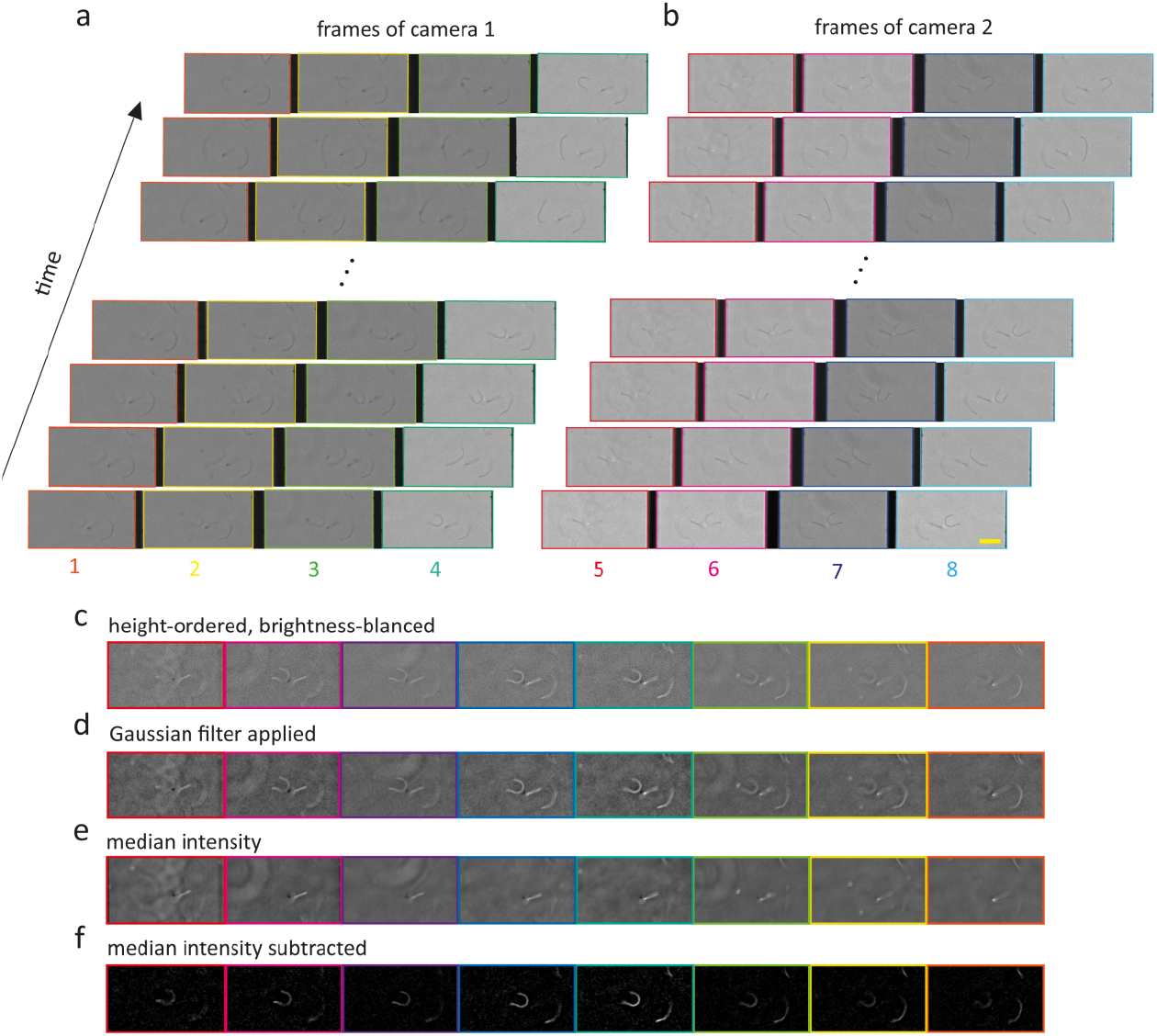
Image acquisition, registering and filtering. (a,b) Four phase contrast images acquired by camera 1 (bottom planes) and camera 2 (top planes), respectively. Length of yellow scale bar in (b) is 10 μm. (c) Image intensities are inverted (for better contrast) and planes are ordered following their distance from the surface, from left (bottom) to right (top). Brightness values of all planes are balanced using the ε(plane #) values from the calibration measurement. (d) Image noise is reduced by applying a 2D Gaussian low pass filter. (e) Median intensity of each plane over 1980 frames. (f) Subtraction of median intensity images for enhancing image contrast.

### Brightness correction, image conversion, and Gaussian filtering

Intensity values in all eight image planes are balanced by multiplying with the corresponding weight factors ε^−1^ (Fig. 1c). All image values are inverted to negative numbers for contrast enhancement and better visualization of axonemes (Fig. 1c). For noise reduction, a 2D Gaussian kernel (`imgaussfilt` function in MATLAB) with standard deviation of σ = 2 is used for low-pass frequency filtering (Fig. 1d).

### Subtraction of median intensity

Median intensity of each plane determined overall frames (Fig. 1 e) is subtracted from data to remove all contributions from immobile and static sources. This also increases the SNR by a factor of 3 to 5 times, depending on signal strength (Fig. 1f).

### Co-registration of planes

The eight raw images have slight relative shear to each other (Fig. 2a) and have to be registered. This is done by calculating the two-dimensional crosscorrelation of each plane to one reference plane (either plane #4 or plane #5 close to the center of the stack) (Fig. 2b). The peak position of this cross-correlation gives direction and value of the relative shift of the considered plane with respect to the reference plane. This shift estimation is refined by repeating the procedure on Fourier up-sampled images, see Stein et al. (2015). Using these values, all images are registered to a common frame with sub-pixel precision (Fig. 2c,d). Non-overlapping image borders arising from registration are cropped in the final aligned image stack (Fig. 2d,e).

**Appendix 2 Figure 2.**
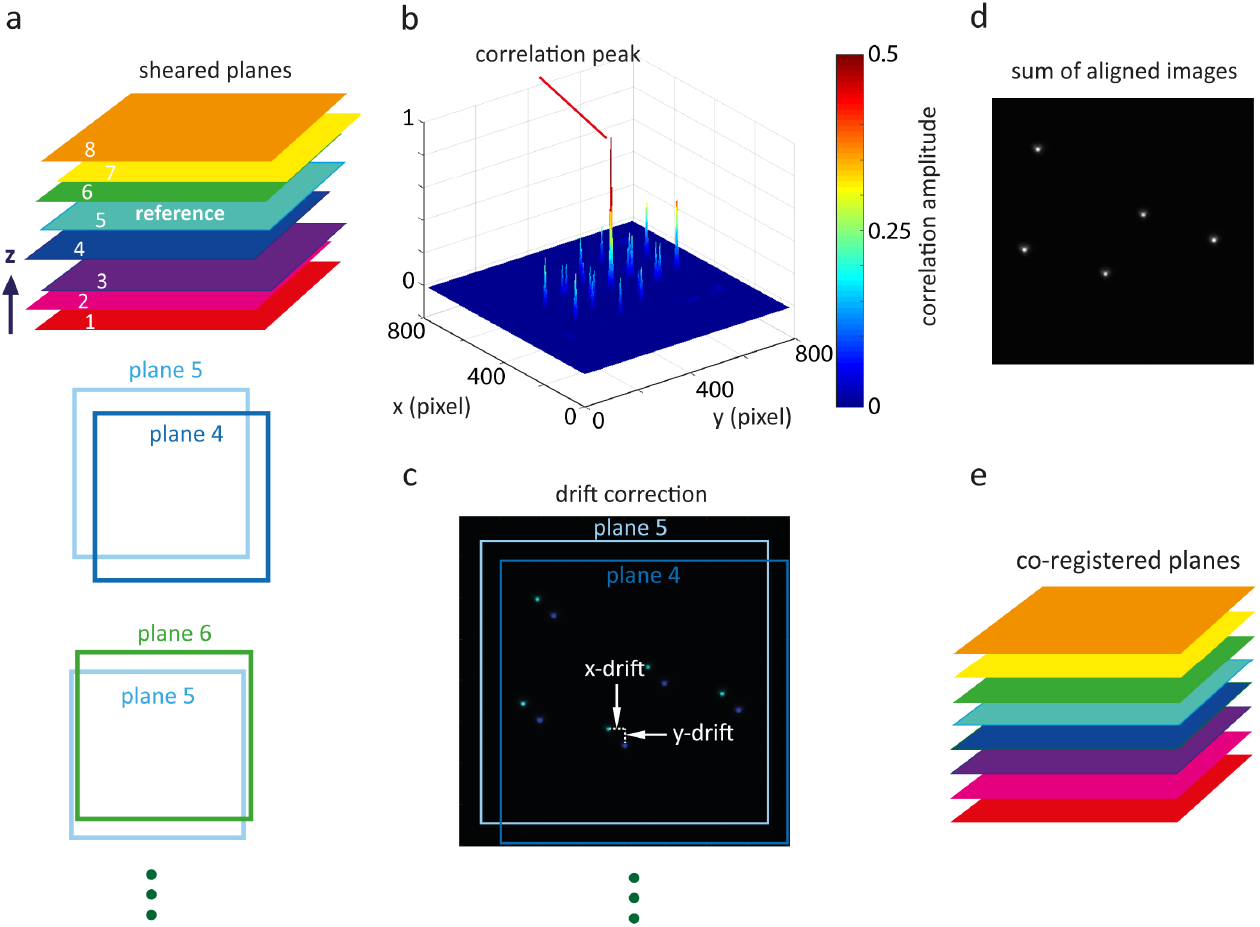
Shear correction. (a) Example of plane shearing where fifth plane from bottom is considered as reference image. (b) Cross-correlation between one plane with reference. Peak position defines direction and value of shear between considered plane and reference plane. (c) Shear correction between plane 4 and 5. (d) Overlay of aligned planes 4 and 5. (e) Shear-corrected image stack of eight planes.

### Image sharpening

Images of *Chlamydomonas* axonemes with diameter of ~ 150 nm (much smaller than the illumination wavelength used). Note that the basal end of the axoneme is slightly (~10%) thicker than its distal tip Geyer (2013), but otherwise all axonemes show an almost homogeneous image intensity along their contour length. Thus, the shape of an axoneme can be effectively considered as a continuous one-dimensional line in three-dimensional space. This *a priori* information can used for a three-dimensional image deconvolution. Each frame of the multi-plane image stack is deconvolved using a pre-calculated aberration-free PSF at the interface of glass coverslide and water (Fig. 3a). This PSF is calculated using the full wave-optical theory developed by Richards and Wolf Wolf (1959); Richards and Wolf (1959) and the optical parameters of our setup (imaging wavelength *λ_peak_* of 585 nm, numerical aperture NA of 1.25, image magnification of 80×, principal focal length of objective of 3 mm, and water and oil refractive index values of *n_water_* = 1.33 and *n_glass_* = 1.52, respectively). Within a small axial distance from the surface, this PSF remains an excellent approximation of the actual ones despite refractive index mismatch, as can be seen by comparing its shape at a nominal focus position of ~1.5 μm above the surface (Fig. 3b) with the aberration-free PSF directly at the surface (Fig. 3a). For deconvolution, we performed ten iterations using a 3D Lucy-Richardson algorithm Richardson (1972); Lucy (1974) (`deconvlucy` function in MATLAB) which results in a sharpening of the axoneme contour by a factor of ~ 2 in all directions. Fig. 3c,e present an axoneme image maximum-projected along the optical axis and along the *y*-direction, respectively. Corresponding deconvolved images are shown in Fig. 3d,f.

**Appendix 2 Figure 3.**
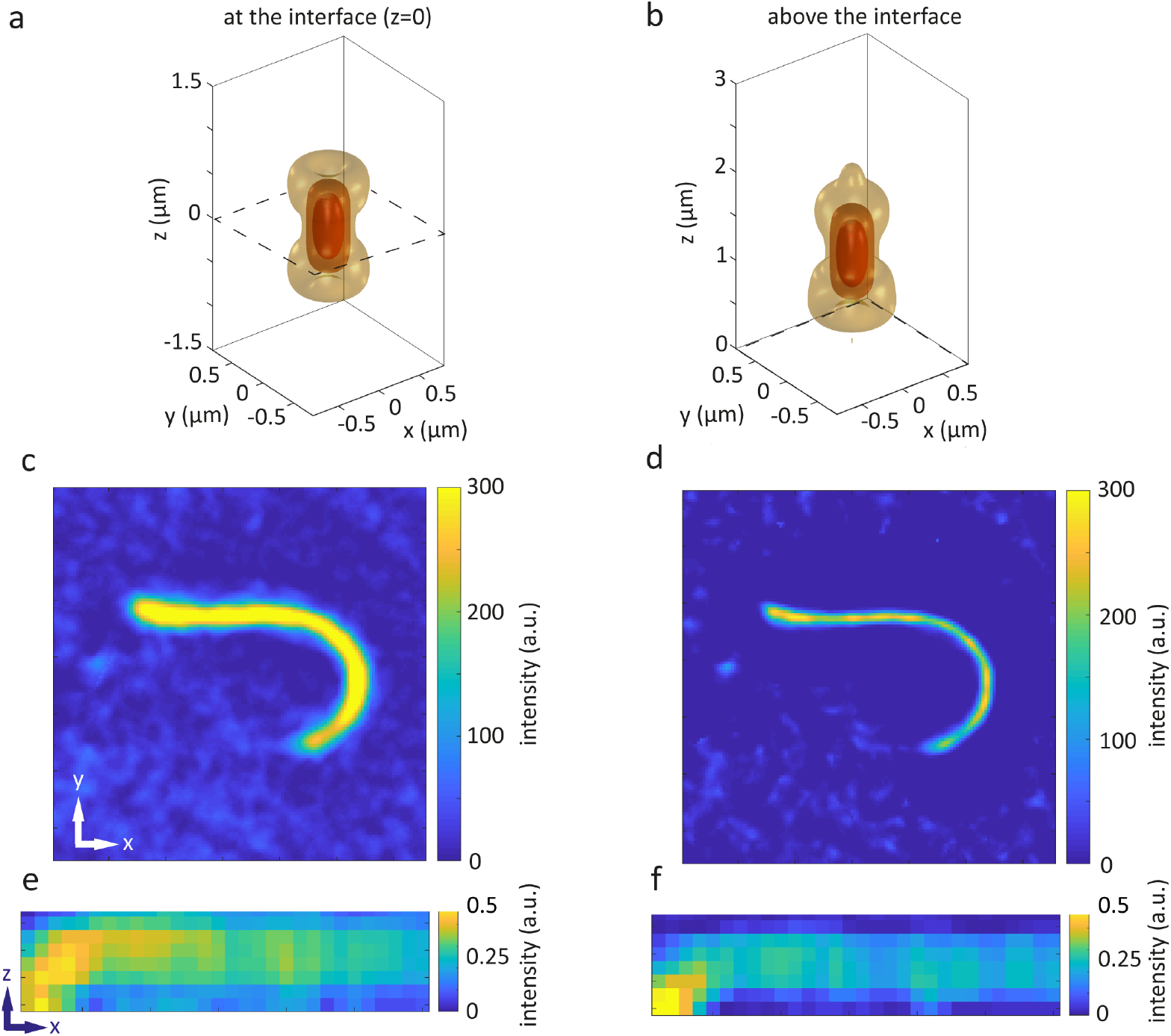
Image sharpening by 3D deconvolution. (a) Iso-surfaces (at 1/*e*, 1/*e*^2^ and 1/*e*^3^ of its maximum value at the center) of a calculated PSF directly at the glass-water interface (indicated by dashed line). The same for a nominal focus position of 1.5 μm above the glass interface. Although small changes can be seen for the 1/*e*^3^ iso-surface, the center shape and values of the PSF are nearly identical to those of the aberration-free PSF in panel (a). (c) Maximum projection of axoneme image along the optical axis. (d) The same image after deconvolution. (f) Maximum projection of axoneme image along a lateral direction. (e) Same image after deconvolution.

## Appendix 3 3D tracking

Tracking of an axoneme is performed by first finding its 2D contour in the 2D maximum intensity projections using a GVF snake Xu and Prince (1997); Xu et al. (1998) (Fig. 1a,b). For the first frame, the user selects a region of interest that should contain only one single axoneme. Then, the user initializes the snake by drawing a polygonal line along the contour of the axoneme (Fig. 1c) which estimates the contour length. In the same frame, the polygonal line is interpolated to *N* = 30 points and used as a starting guess for the snake algorithm (Fig. 1d). The GVF is calculated using 20 iterations and a GVF regularization coefficient of μ = 0.1. The snake is then deformed to the contour in the next frame according to the GVF where we have adapted the original algorithm by Xu and Prince for open boundary conditions Xu et al. (1998).

In a next step, a height profile *h_i_*(*z*) is created for each segment *i*, averaging the intensity over pixels of the segment (Fig. 1e). This profile is fitted with a 1D Gaussian (Fig. 1f). The *z*-position of the maximum of this fit yield the axial coordinate *z*, of the segment, and together its lateral positions *x_i_* and *y_i_* constitute the 3D coordinates of segment *i*.

**Appendix 3 Figure 1.**
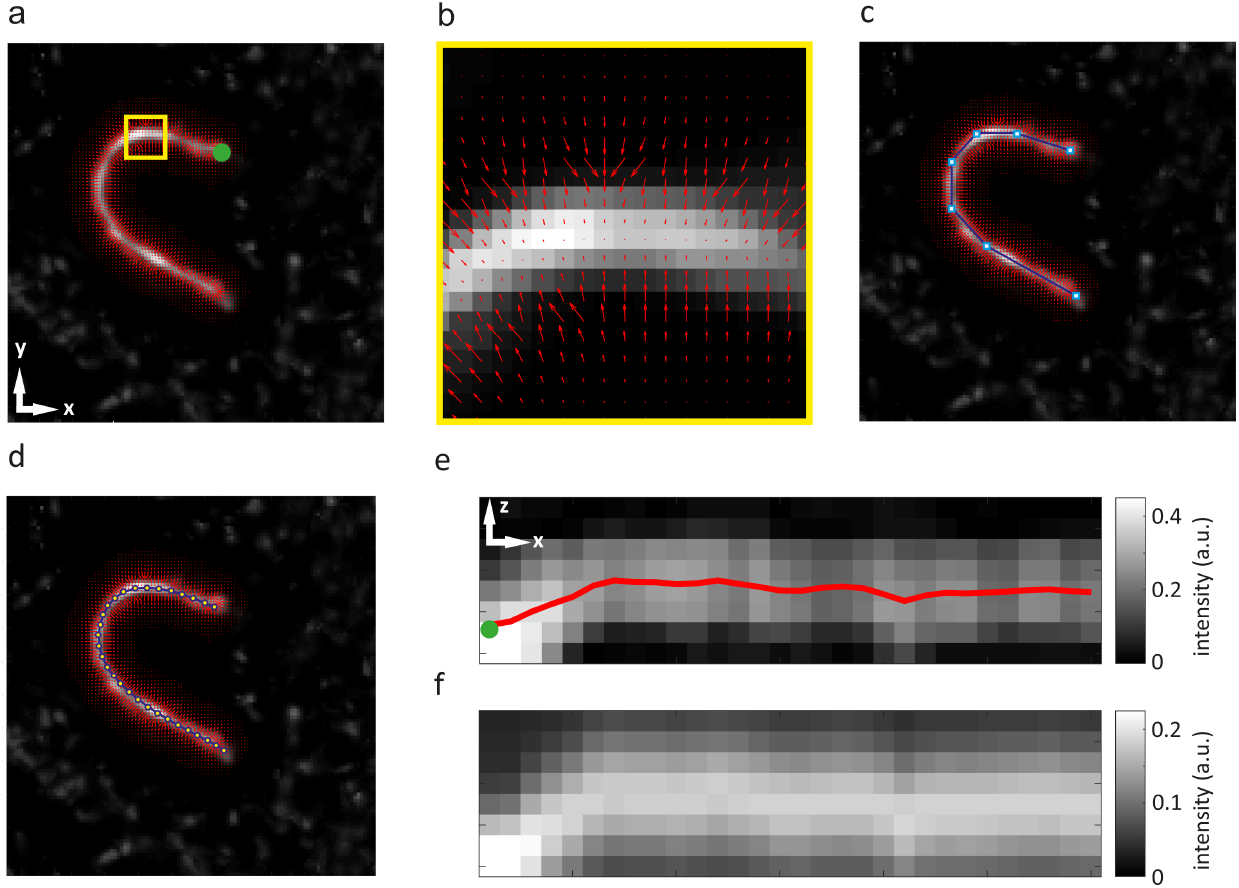
3D tracking of axonemal motion. (a) Maximum intensity projection of axoneme image (green dot indicates basal end). (b) Calculated vector for small yellow region of panel (a), converging toward pixels with maximum intensity. (c) A polygonal line with seven nodes along the contour snake. (d) Division of snake into 30 segments. (e) Vertical cross section of axonemal image and contour (green dot indicates basal end). (f) Same as (e), but after fitting axial intensity distribution at each lateral position with a 1D Gaussian.

An installation file ‘AxonemeTracking3D.mlappinstall’ for a platform-independent MATLAB app together with its source code (‘source code.zip’) and sample data (‘testdata.mat’) can be downloaded at: https://www.dropbox.com/s/x8iequg72zmkx4b/FlagellumAnalysis.zip?dl=0. This app executes the contour determination and contour discretization of 3D axoneme images. The zip-file contains also the commented MATLAB program ‘FlagellumAnalysisExample.m’ together with tracking data (‘result_tracking22.mat’) of the flagellum analyzed for Fig. 3a,b which demonstrates the calculation of curvature and torsion from a discretized contour, and their graphical display.

## Appendix 4 Contour fitting

As described in the main text, contour fitting of the three-dimensional contour obtained from the 3D tracking procedure was done by a polynomial fit of the determined discrete 3D coordinates. Such a fit become ambiguous at the ends of the tracked contour which can lead to unphysically large values of curvature at these ends. To prevent this, we have padded a discretized contour on both ends by mirrored inner points. The impact of the number of padded points on the fit results is shown in Fig. 1. As can be seen, the absolute values of curvature at the contour ends become smaller with increasing contour padding, but the values in the middle of the contours remain mostly unchanged, similar to the torsion values.

**Appendix 4 Figure 1.**
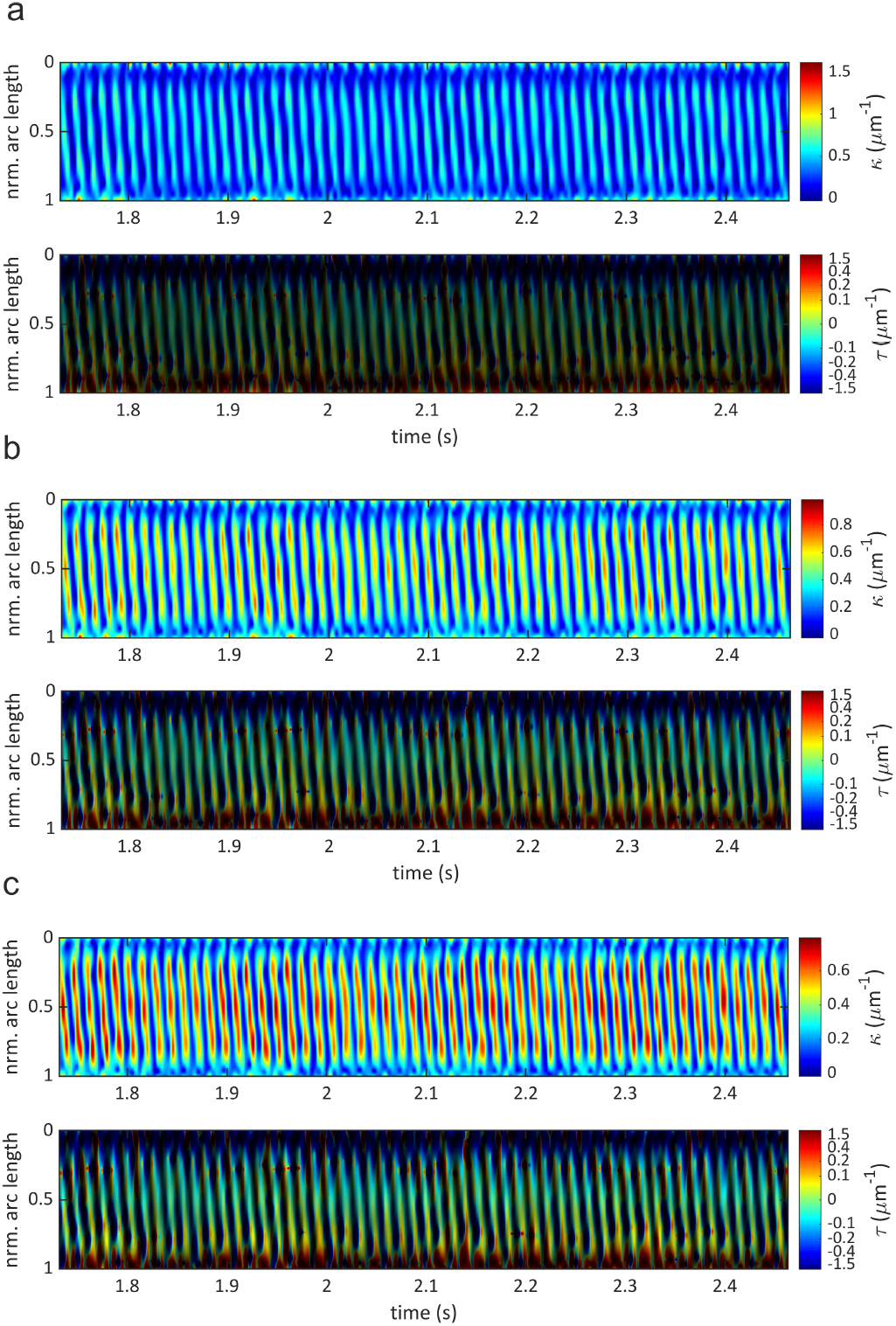
Effect of contour padding on calculated curvature and torsion. (a) Zero padding, (b) padding with one extra point, (c) padding with two extra points. Compare with Fig. 3 in main text showing results for padding with three extra points. As can be seen, padding does mostly suppress excessive curvature values at contour ends, but does nearly not change curvature values elsewhere, and has little impact on torsion values (different brightness values in torsion plots are due to different dynamic range of curvature values).

Using the results of the polynomial fits, we estimated the fit accuracy (and checked for any systematic fit bias) by histograming the difference between the discrete coordinates as delivered by the axoneme tracing algorithm and the positions calculated form the polynomial fit. The resulting histograms for both axonemes analyzed in Fig. 3 are shown in Fig. 2. The symmetry of the residual histograms indicates that there is no systematic bias of the polynomial fit that would lead to skewed asymmetric distributions of the residuals. The width of the distributions gives an estimate of the accuracy of the contour position determination. The standard deviations σfor each distribution are indicated above each histogram. We attribute the fluctuations of the discretized positions (as delivered by the tracing algorithm) around the smooth polynomial fit to noise-induced uncertainties of the contour tracing.

**Appendix 4 Figure 2.**
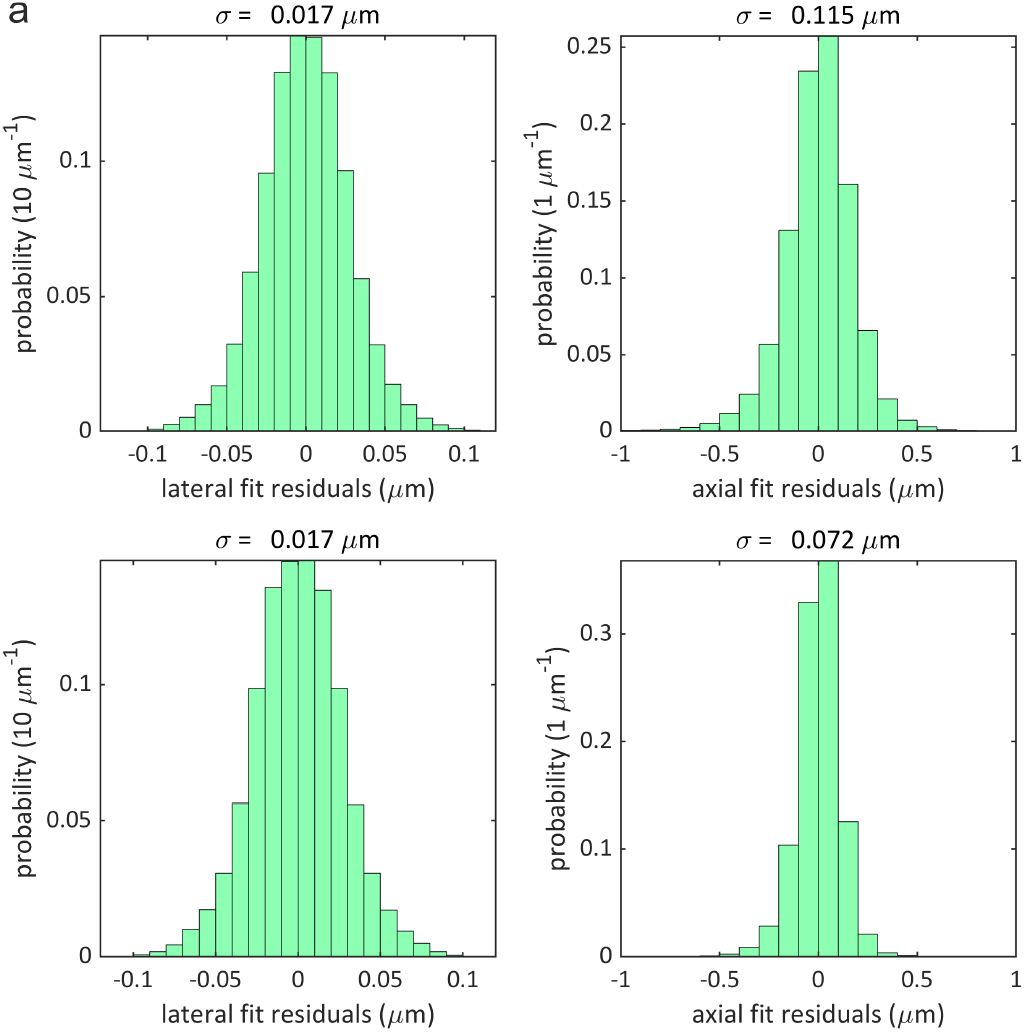
Distributions of residuals between discretized and fitted position values for (a) flagellum of Fig. 3a,b and (b) for flagellum of Fig. 3c,d. Left panels show residuals for lateral *x,y*-positions, right panels for axial *z*-positions. Calculated standard deviation values for all distributions are given on top of each panel.

## Appendix 5 List of hardware components used in multi-plane phase contrast microscope

**Appendix 5 Table 1.** Specifications of components incorporated into the multi plane phase contrast microscope

List of hardware components used in multi-plane phase contrast microscope
| Component | Details |
| --- | --- |
| white light source | 100 W 12 V Halogen Lamp U-LH100L-3, Olympus, peak wavelength ~585 nm |
| phase annulus | IX-PH3, 35 mm diameter, Olympus |
| condenser lens | IX2-LWUCD Universal Condenser, Olympus, NA 0.55, WD 27 mm |
| microscope stage | ES107IX2, Prior scientific instruments, 115 × 77 mm travel range, 1 μm step size resolution, OptiScan ES107 Controller, |
| phase objective | UPLFLN 60XOIPH, 0.65-1.25 NA, Olympus |
| tube lens | $f_{TL} = 180$ mm, Olympus |
| field aperture | SP 40 slit, Owis |
| telescope lenses | $f_{L1} = 150$ mm, $f_{L2} = 200$ mm, achromatic doublets, AC254-150-A-ML, AC254-200-A-ML, Thorlabs |
| prism | Corning C-7980, $n_D = 1.458$, Abbe number $v = 67.8$, see section 6.2 in SI of ref. *Descloux et al. (2018)* |
| camera | ORCA-Flash 4.0 V2, Hamamatsu, pixel size = 6.5 μm |

